# Microbial community structure is affected by phage-resistance associated increases in host density

**DOI:** 10.1101/2024.03.07.583923

**Authors:** Meaghan Castledine, Daniel Padfield, Rai Lewis, Angus Buckling

## Abstract

Lytic bacteriophages (‘phages’) can limit bacterial densities and shape community structure, either directly through lysis or indirectly through costs to resistance. However, phages have also been reported to have no, and in some cases even positive, effects on host densities. Here, we investigate the mechanisms behind an increase in host density in *Variovorax* sp. populations following a fixation of resistance that was maintained after phage extinction. Our results demonstrate that the density increase was a genetic trait coinciding with resistance emergence. Growth curves showed that phage resistance shifted population growth curves such that density was higher in the death-phase. This density-increasing effect of resistance had important implications for community structure with phage resistant *Variovorax* decreasing the density of a conspecific. That resistance to lytic phage can increase host densities has implications for wider ecology and phage therapy where lytic phages are presumed to have negative effects on their hosts.

## Introduction

Bacteriophages are hypothesised to have major impacts on structuring microbial communities, and in regulation of wider nutrient cycles (Winter *et al*. 2010; Breitbart 2012; Breitbart *et al*. 2018). Part of this assumption comes from the ubiquity of bacteriophages (Clokie *et al*. 2011). By lysing their hosts, phages can suppress fast-growing bacteria densities and give a relative advantage to slow-growing or rarer microbial taxa (Winter *et al*. 2010). Phages are shown to lyse algal blooms, resulting in crashes in algae abundances and nutrient release which benefits the surrounding community (Breitbart *et al*. 2018; Flynn *et al*. 2022; Liao *et al*. 2023). Furthermore, a common assumption of many models is that phage resistance is costly (Winter *et al*. 2010), which leads to suppression of dominant or previously-fast growing bacteria populations (Gandon *et al*. 2008; Winter *et al*. 2010).

However, the effects of phage resistance on host phenotypes, and the associated cost, is highly system specific. If evolving phage resistance typically involves modification of a receptor essential to host growth, resistance is likely to be associated with a growth rate cost (van Houte, Buckling and Westra 2016). For example, to become resistant to OMKO1, *Pseudomonas aeruginosa* mutates an efflux pump required for drug resistance (Chan *et al*. 2016). As a result, while antibiotics are present, resistance to OMKO1 imposes a cost to *P. aeruginosa* (Chan *et al*. 2016). Similarly, phage resistance involving mutations to flagella or pili are further shown to slow bacteria growth rates (Buckling *et al*. 2006; Li *et al*. 2022). However, there are numerous examples of phage resistance being non costly to growth including in *Pseudomonas syringae* (Koskella *et al*. 2012), *Synechococcus* (Lennon *et al*. 2007), and *P. aeruginosa* (Castledine *et al*. 2022).

Phage resistance may also indirectly increase host fitness (beyond the direct effect of resisting phage). For instance, increased biofilm production is a phage resistance mechanism which physically blocks phage access to bacteria cells (Salathé and Soyer 2008; Harper *et al*. 2014; Abedon 2023). Biofilm production has consequential fitness benefits in resisting wider environmental stress such as antibiotics (Crabbé *et al*. 2019), and can be advantageous in competition (Oliveira *et al*. 2015). Phages can also accelerate the molecular evolution of their hosts and select for hypermutator strains, therefore giving increased adaptive potential to secondary stressors (Pal *et al*. 2007; Jayaraman 2011). *P. fluorescens* phage has been shown to increase host densities in soil, although only when grown in monoculture (in the absence of a soil community) (Gómez and Buckling 2011). Here, phages are hypothesised to have increased bacterial densities by reducing intraspecific competition through slowing of growth rates or facilitating diversification into separate ecotypes (Gómez and Buckling 2011), although the exact mechanism is unknown.

Understanding how phages can increase bacteria densities is important for phage therapy where increasing bacteria densities would be undesirable (Abedon 2023). Furthermore, investigating this effect has important implications for our understanding of how phages shape microbial communities where, in nature, phages are presumed to have predominantly negative effects their host densities (1, 3). In a previous study, we noted that densities of *Variovorax* sp. increased following phage exposure and fixation of phage resistance (Castledine *et al*. 2024a) even when phage had been driven extinct. Here, we investigate the mechanism underpinning phage-mediated increases in *Variovorax* densities, and the consequence this has for the composition of a three species synthetic bacteria community. We show that phage resistance is associated with density increases resulting from a shift in growth curves that led to higher densities in the death-phase. Phage resistance had ramifications for community structure with decreases in the proportion of another community member which is parasitised by *Variovorax*.

## Materials and Methods

### Experimental evolution

In a previous study, species *Ochrobactrum* sp., *Pseudomonas* sp., and *Variovorax* sp. were grown as monocultures and as a three-species polyculture with and without their respective phage (*Ochrobactrum* phage ORM_20, *Pseudomonas* phage CH7FMC, *Variovorax* phage VAC_51) (Castledine *et al*. 2024a). Experimental treatments included: monoculture, monoculture with phage (each bacterial species individually with only its species-specific bacteriophage), polyculture (all three bacteria present) and polyculture with phage (all three bacteria with all three phages present) (Figure 1). Each treatment was replicated six times. Experimental set-up is described in (Castledine *et al*. 2024a). Briefly, cultures were established using isogenic strains of each bacteria and phage, with phage added at an MOI of 0.01 (to phage present treatments). Cultures were grown in 1/64 TSB (tryptic soy broth diluted in demineralised H_2_O), at 28 °C and transferred weekly to fresh medium. Bacteria and phage densities were estimated every two weeks. For *Variovorax* cultures, we estimated phage resistance in week two and eight monocultures (phage present and absent). Phage resistance was analysed using spot assays with ancestral phage.

**Figure 1.**
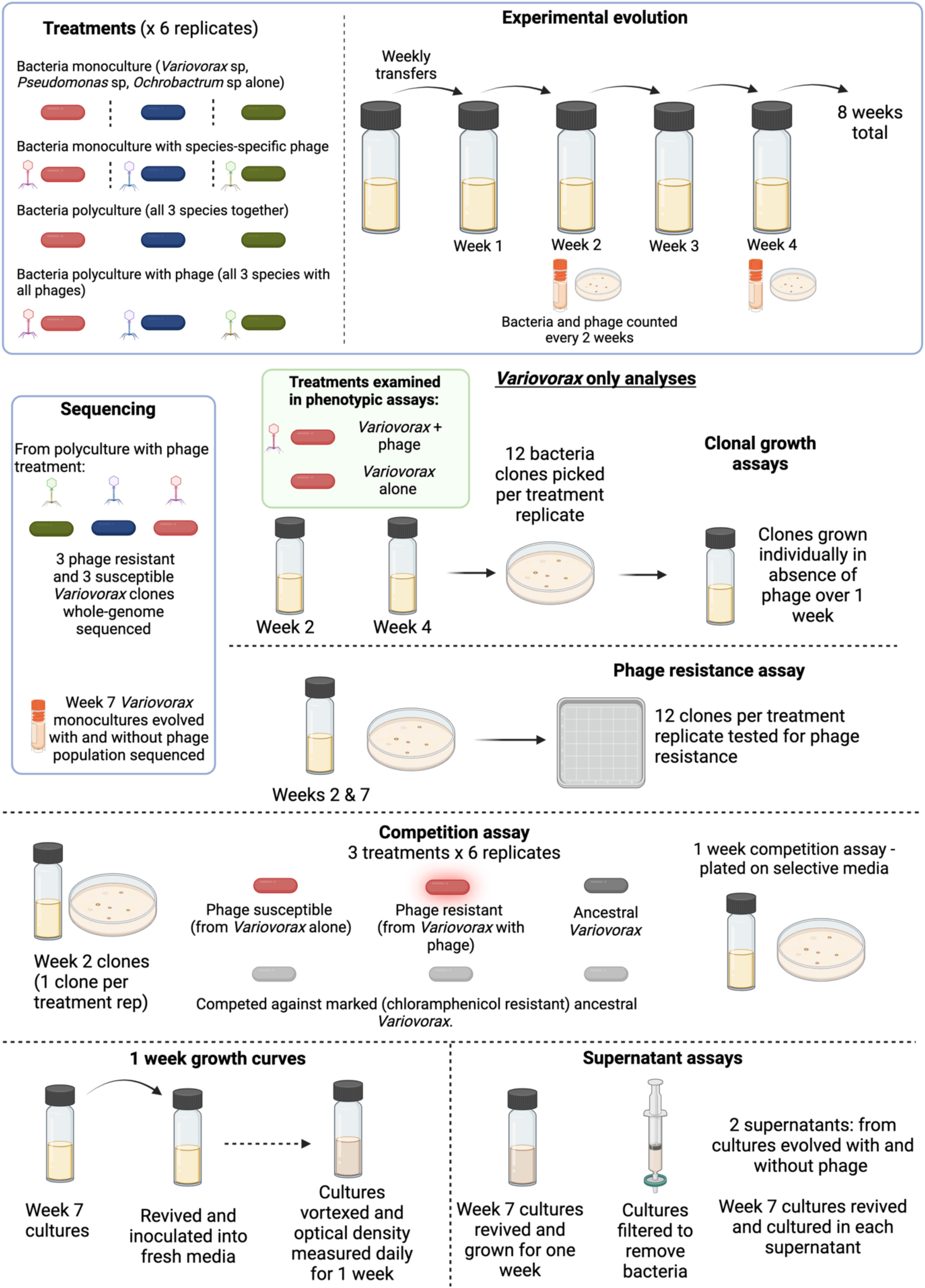
Overview of experimental design in which *Variovorax* and its phage evolved in monoculture and polyculture, and assays performed to understand the mechanism of this density increase. Sequencing analyses and phenotypic assays aimed to determine the mechanism by which phages led to increased *Variovorax* densities across experimental evolution. Created in BioRender. Castledine, M. (2024) <u>BioRender.com/c73o264</u>.

### Clonal density measures

We first wanted to assess whether higher densities conferred to phage-exposed populations was a heritable trait observable in single clones picked and regrown in the absence of phage. From week two and four cultures (the weeks preceding and following higher densities), twelve colonies were isolated from each replicate from monoculture treatments. Colonies were individually grown in 1/64 TSB and density normalised as before. To fresh 6 mL 1/64 TSB, ∼10^6^ CFUs of each clone were added to individual vials. Cultures were grown for 1 week, frozen, and plated from frozen onto KB agar as previous.

### Clonal and population sequencing and bioinformatics

To gain insight into the mechanism of resistance and whether this could be linked to changes in bacterial density and growth rate, isolates were sent for whole genome sequencing (WGS). Three phage resistant and three phage susceptible clones from three independent week two polyculture lines were selected for WGS. Polyculture lines were selected as these were the only treatment which had phage susceptible and resistant clones coexisting (100% resistance was found in monoculture lines). This allowed for direct comparison between different clones that have evolved from the same starting population but only differed in their resistance profile. DNA extractions were done on each isolate by MicrobesNG using in-house protocols (MicrobesNG Sequencing Service Methods, 2021). WGS was performed by MicrobesNG using an Illumina Novaseq 6000 to create 250 base pair (bp) paired-end reads. In-house processing by MicrobesNG trimmed the adapters and removed reads with a quality <15.

To further understand genetic changes that resulted in increased growth rates of phage resistant *Variovorax*, cultures from week 7 monoculture lines of evolution were sent for population sequencing. Overnight cultures (120 µL frozen stock in 1/64 TSB grown shaking at 28 °C. 1.8 mL was spun at 16000 x g for 5 mins (Progen GenFuge 24D centrifuge). The supernatant was removed and another 1.8 mL of fresh sample was added to each tube and centrifuged for a further 5 mins and supernatant removed. The Qiagen DNeasy Ultra Clean DNA Extraction Kit reagents and protocol were used to extract the gDNA from the cultures. The concentration of gDNA in each sample was determined using the QuBit dsDNA HS Assay kit regents and protocol. Sequencing was done by the University of Liverpool’s Centre for Genomic Research (CGR) using an Illumina Novaseq to create 150bp paired end reads. The CGR trimmed for the presence of Illumina adapter sequences using Cutadapt (v1.2.1) and removed reads with a minimum quality score of 20 and which were shorter 15bp in length.

For both clonal and population sequencing, we further filtered the trimmed reads to remove Illumina adapter sequences and for reads with a quality score below 10 using TrimGalore (v0.6.4_dev). We used the breseq pipeline (v0.36) – a computational tool for analysing short-read DNA data - to map the trimmed reads to a high quality reference genome of *Variovorax* sp. (GenBank accession: GCA_037482385.1) and predict genetic variants (Deatherage and Barrick 2014). As our reference genome has two contigs, we added the parameters “-c” and “—contig-reference” so that all the sequences are assumed to have the same average coverage and coverage distribution. In the clonal sequencing, 97.6% of reads mapped to the reference genome (minimum of 95.4%), giving on average ∼876,989 reads per replicate (minimum 444,292) with an average coverage of 62.21 (minimum of 31.84). We only investigated genetic variants that were at a frequency of 1 (we do not expect polymorphisms in any of the predicted genetic variants). For the population sequencing, we ran breseq in polymorphism mode (-p). Variants with <5% frequency in the population samples were removed.

### Phage resistance and competition assay

We determined if phage resistance conferred a growth rate benefit to *Variovorax*, thereby leading to higher bacterial densities. Twelve clones per replicate from week two monoculture-lines were isolated and grown in 96-well plates with 150 µL TSB. Phage resistance was determined by streaking isolated colonies against the ancestral phage (streak assay). Week two isolates were chosen as this time-point preceded changes in bacterial density from phage exposure and phage extinction. We elected for phage resistant clones generated during experimental evolution (instead of selecting spontaneous mutants) in the hope that resistance mutations/mechanisms (and therefore phenotypic effects) were consistent with those from the evolution experiments. One phage resistant clone from the phage resistance assays was chosen for each replicate (n = 6), and one clone from each no-phage treatment replicate was isolated as a phage susceptible control (n = 6). Each colony was grown individually in 1/64 TSB individually as described previously. In addition, marked (ancestor-cp; chloramphenicol resistance (Sünderhauf *et al*. 2023)) and unmarked ancestral *Variovorax* sp. were grown. Cultures were diluted to normalise densities as described previously. The unmarked ancestral strain and each phage resistant and susceptible colony was competed against the marked ancestral strain at equal starting densities (∼3 x 10^6^ CFUs of each competitor inoculated (30 µL diluted culture)). Diluted cultures used in inoculation were frozen and plated on KB agar as above for starting density estimation. Cultures were grown for one week as previous and plated from frozen onto KB and KB-chloramphenicol (25 μg/mL) agar. Relative fitness was calculated from the ratio of the estimated Malthusian parameters, *m*_resistant_, *m*_susceptible_ or *m*_ancestor_:*m*_ancestor-cp_, which were calculated as *m* = ln(*N*_1_/*N*_0_), where *N*_1_ is the final density and *N*_0_ is the starting density (Lenski *et al*. 1991).

### Growth curves

Next, we were interested if phage resistance had shifted the growth curve of *Variovorax* populations thereby leading to higher densities at transfer. 120 µL from each monoculture line from week seven were revived in 6 mL growth medium and grown shaking for two days at 28 °C. Densities were normalised as previous and 10 µL was inoculated into fresh growth medium. Every 24 hrs for one week, cultures were vortexed and optical density (OD_600_; Multiskan Sky) was measured from a 200 µL sample.

To characterise the log/exponential growth phase of phage resistant and susceptible cultures at a higher resolution, week seven cultures were revived as previous and 20 µL of diluted culture (∼10^6^ CFUs) were inoculated into 180 µL growth medium. Optical density reads were measured every 45 mins for 24 hrs.

### Supernatant assay

We hypothesised that higher densities and shifts in growth curve may be caused by production of less-toxic metabolites. This hypothesis was tested using a supernatant assay. Week seven cultures were revived and normalised as previous using five replicates of resistant and susceptible populations (one replicate each of susceptible and resistant cultures became contaminated and were not used in the experiment). Two replicates of each week seven culture (two technical replicates for each biological replicate) were established to ensure enough supernatant was available for the reciprocal transplant assay. Cultures were grown for one week. On day five, fresh week seven cultures were revived and grown for two days. Week-old cultures were sterilised by passing medium through a 0.22 µm filter syringe. Replicate supernatants from phage resistant and susceptible populations were pooled into separate supernatant stocks. To separate vials, 3 mL of phage resistant and susceptible supernatant was added to 25 mL vials. Two-day (fresh) culture densities were normalised and each culture was inoculated into both supernatant types. This established two controls in which cultures were grown in the same supernatant (e.g. phage resistant in phage resistant supernatant) and treatments in which cultures were transplanted into the opposite supernatant (e.g. phage resistant in phage susceptible supernatant and vice versa). Cultures were grown for two days and plated as previous.

### Phage resistance and community structure

Next we assayed whether resistance to phage gives *Variovorax* an advantage over *Ochrobactrum* and *Pseudomonas* when phages are absent. Six independent phage resistant clones and six independent phage susceptible clones were isolated from phage and no phage week two monoculture evolution lines respectively. ‘No phage’ lines were selected for isolation of phage susceptible clones as phage resistance was too high in ‘phage’ lines for enough individual susceptible clones to be isolated. Comparison between ‘no phage’ and ‘phage’ lines accounts for any change in general laboratory adaptation. Each clone and ancestral *Variovorax* were individually grown for two days at 28 °C, shaking (180 r.p.m.) in 6 mL 1/64 TSB. Ancestral *Pseudomonas* and *Ochrobactrum* isolates were also grown in the same conditions. Isolate densities were normalised to 10^5^ CFUs/mL and 10 μL added to relevant vials. Each *Variovorax* clone (ancestral, resistant, susceptible) was grown as a polyculture with *Pseudomonas* and *Ochrobactrum*, with six replicates per treatment. Cultures were grown static for 1 week at 28 °C, then frozen as previous and plated from frozen.

### Statistical analyses

All data were analysed using R (v.4.2.1) in RStudio (Team 2013) and all plots were made using the package ‘*ggplot2*’ (Wickham 2016). Model simplification was conducted using likelihood ratio tests and Tukey’s post hoc multiple comparison tests were done using the R package ‘*emmeans*’ (Lenth 2018).

*Variovorax* densities were analysed in a linear mixed effects model with density (log10 CFU/mL) tested against interacting fixed effects of phage exposure, culture type (polyculture or monoculture) and time (weeks two, four, six, eight) with a random effect of treatment replicate. Clonal densities (log10 CFU/mL) from clonal growth experiments were analysed against interacting fixed effects of phage exposure and time (week two or four) with random effects of treatment replicate and block (experiment conducted in two blocks for feasibility). In competition assays, relative fitness was analysed in a linear model with a fixed effect of competitor (ancestor, phage resistant or susceptible).

One week growth curves were analysed in a mixed effects model with optical density tested against a fixed term of day and interacting fixed effects of treatment (phage resistant / susceptible) and a quadratic term of day to account for the non-linear relationship. A random effect of treatment replicate was included. 24-hour assays of bacterial growth were used to estimate exponential growth rate. One replicate was removed from the no-phage treatment for failing to grow. Per replicate, exponential growth was estimated using rolling regression, taking the steepest slope of the linear regression between *OD*_600_ and time in hours in a shifting window of every nine time points (∼6.75 hr). A linear model tested growth rate against treatment.

The supernatant assay was analysed in a linear mixed effects model with density (log10 CFU/mL) analysed against interacting fixed effects of supernatant type and original treatment with a random effect of treatment replicate.

To test whether consistent genetic differences occur within treatments, we performed nonmetric multidimensional scaling on the Euclidean distance matrix of SNPs/indels and their proportions in each population using ‘*metaMDS*’ in the R package ‘*vegan*’ (Oksanen *et al*. 2019). Nonmetric multidimensional scaling aims to collapse information from multiple dimensions (i.e., from different populations and multiple SNPs/indels per population) into just a few, allowing differences between samples to be visualized and interpreted. Permutational ANOVA tests were run using ‘*vegan::adonis*’, with Euclidean distance as the response term and treatment as the predictor variable. Additionally, we analysed whether evolution with phage affected the genetic difference of each population (phage, no phage) from the ancestral population. The genetic distance from the ancestral population was calculated as the sum of the difference of the proportion of each SNP/indel in each population from the ancestral proportion. Genetic distance was then analysed in a linear model analysing distance against phage presence/absence.

To assess whether *Variovorax* phage resistance influences community structure in the absence of phage, we analysed the relative proportion of each species in a generalised linear mixed effects model. Here, each species proportion was analysed against interacting fixed effects of *Variovorax* phage resistance and species identity, with a random effect of treatment replicate and a binomial error structure.

## Results

### Phage resistant clones have higher densities than phage susceptible clones

In a previous study (Castledine *et al*. 2024a), we evolved *Variovorax* and its lytic phage in the presence and absence of two other bacteria-phage pairs (Figure 1). We observed an interesting interaction with *Variovorax* densities increasing after two weeks of phage exposure (phage present: 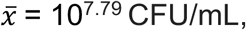 95%CI = 10^7.74^ - 10^7.85^; phage absent: 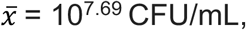 95%CI = 10^7.63^ - 10^7.74^; ANOVA comparing models with and without phage: 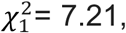 p = 0.007; Tukey HSD: p = 0.013). This coincided with a fixation of phage resistance in polyculture (90.3%; SE ±6.6) and monoculture (100%), and extinction of phage after two weeks. As such, we investigated the hypothesis that phage resistance has led to increased *Variovorax* densities.

To assess whether higher densities in phage-exposed populations was likely a result of a genetic change (as opposed to epigenetic or physiological), we measured the densities of individual clones (isolated from monoculture treatment on weeks two (pre phage loss) and four (post phage loss)) grown in isolation (no phage or other clones present so any density effects are purely heritable). All clones isolated from phage treatments were phage resistant as phage resistance reached fixation in monoculture, while all clones from the no-phage treatment were susceptible. Consistent with the population dynamics during experimental evolution, clones isolated from phage-exposed replicates had 66% higher densities (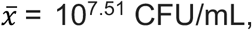 95%CI = 10^5.60^ - 10^9.41^) than phage-unexposed clones (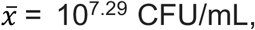 95%CI = 10^5.38^ - 10^9.19^; ANOVA comparing models with and without phage: 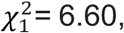 p = 0.0102; Tukey HSD: p = 0.018; Figure 2). Additionally, there was a significant independent effect of time with clones from week four (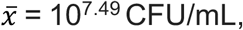 95%CI = 10^5.30^ - 10^9.68^) reaching higher densities than clones from week two (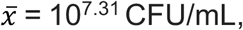 95%CI = 10^5.12^ - 10^9.49^; Tukey HSD: p < 0.001; Figure 2). Overall, these results suggest effects imposed by phage were hereditary as phage did not need to be actively present for effects to be observed.

**Figure 2.**
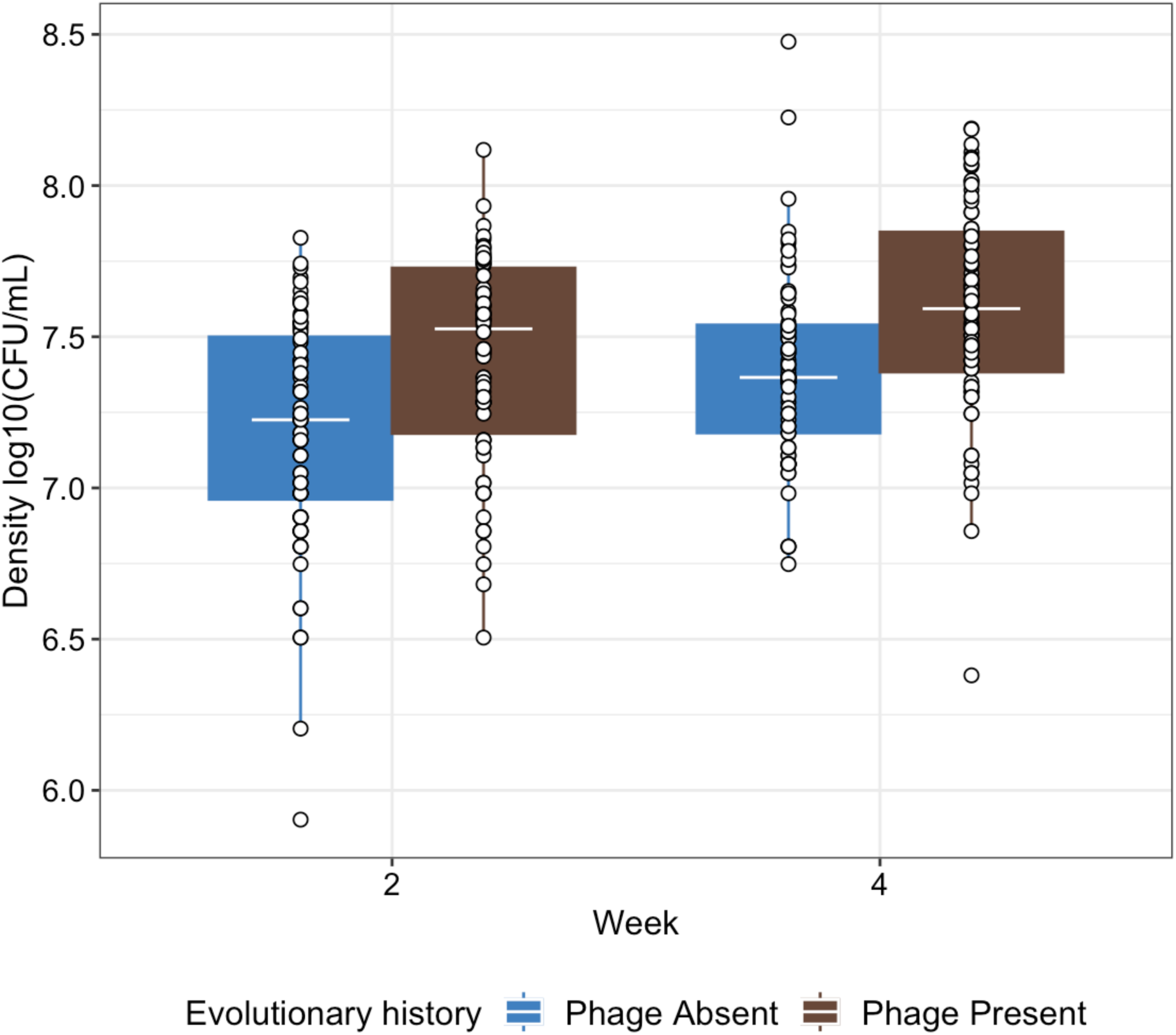
The density of individual clones (points) which have been grown after being isolated from evolution lines where phage have been absent or present. Tops and bottoms of the bars represent the 75th and 25th percentiles of the data, the middle lines are the medians, and the whiskers extend from their respective hinge to the smallest or largest value no further than 1.5* interquartile range. Points represent individual clones.

### Phage resistant populations have distinct mutations

Given the consistent higher growth rate that phage exposed populations and phage resistant clones had over phage absent populations and phage susceptible clones, we looked at whether there were underlying genetic changes that could explain this. We sequenced 3 resistant and 3 sensitive clones coexisting within 3 polyculture treatment replicates, allowing for a direct comparison between different clones that evolved from the same starting population but differed in their resistance profile. Of the phage resistant clones, 2/3 clones had a missense mutation at the same locus (gene ID: 01575) and 1 had a substitute mutation at a nearby locus (gene ID: 01584; 10.4k base pairs downstream; this was also the only mutation in this clone) that were not found in phage susceptible isolates. No other mutations were found at the same loci across clones. Single point mutations are consistent with surface modification forms of resistance (Beckett and Williams 2013; van Houte, Buckling and Westra 2016). Generally, few mutations were found, with one resistant clone having four mutations while the other two had only one mutation each. Similarly, one phage susceptible clone had two mutations while the other two had one mutation. 01584 sequence (mutation found in 1/3 resistant clones) had no comparable genes in BLAST, however its proximity to the other genes could imply a similar function (Demerec and Hartman 1959; Ballouz *et al*. 2010).

We conducted pool sequencing analysis of seven weeks of evolution in monoculture (*Pseudomonas* and *Ochrobactrum* absent), with the aim of identifying further genetic changes linked to phage resistance and increased density. 5/6 phage resistant populations had at least one genetic variant at high frequency (>50%) that was not present in the phage susceptible populations, compared to only 2/6 of the phage susceptible populations, suggesting specific mutations were selected by phage (Figure S1). However, a lack of convergence suggests phage resistance may be maintained by several different mechanisms that have the same phenotypic effects (resistance and higher population density) (see Supplementary Results for full analysis).

### Phage resistant clones do not have a fitness advantage

We next determined whether phage resistance conferred a fitness advantage in the absence of phage. In competition assays with a marked ancestral strain, relative fitness was not significantly affected by phage resistance (F_2,15_ = 0.460, p = 0.640; Figure 3). The greater density associated with resistance did not therefore result in a competitive fitness advantage.

**Figure 3.**
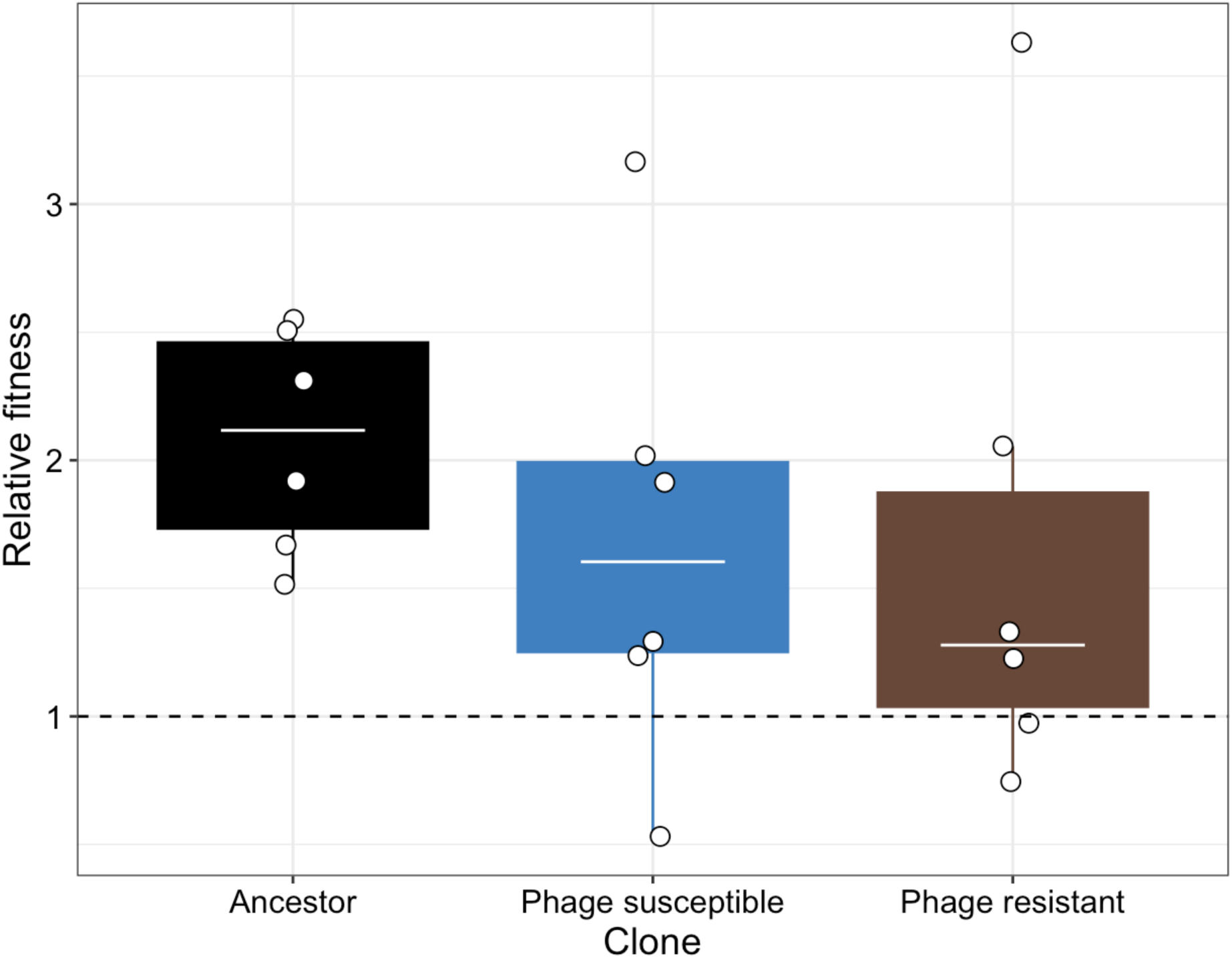
Relative fitness of ancestral, phage susceptible and resistant isolates against a marked ancestral strain of *Variovorax*. Points represent independent replicates. Tops and bottoms of the bars represent the 75th and 25th percentiles of the data, the middle lines are the medians, and the whiskers extend from their respective hinge to the smallest or largest value no further than 1.5* interquartile range.

### Phage resistant populations have greater persistence during the death-phase

We next investigated if density benefits resulting from resistance arose from changes in growth dynamics. This may have arisen if, for example, logistic growth rates of resistant isolates were slowed, resulting in resources becoming less exhausted when cultures were transferred to fresh media on day seven. As such, we measured the growth curves of *Variovorax* populations isolated from week seven of experimental evolution over one week. Here, there was a significant interaction between treatment and day of measurement (ANOVA comparing models with and without the interaction: 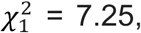 p = 0.007), suggesting phage resistance had altered population growth dynamics (Figure 4). Consistent with our previous observation based on CFU counts (Figure 2), optical density after one week was higher in phage resistant (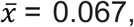 95%CI = 0.062 - 0.072) than susceptible (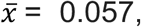 95%CI = 0.053 - 0.062) populations (Figure 4). Consistent with an absence of fitness differences between susceptible and resistant isolates (Figure 3), there was no significant difference in exponential growth rate measured in the 24 hr growth rate assay (F_1,9_ = 0.622, p = 0.451) or carrying capacity in the one week growth rate assay (phage resistant: 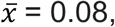 95%CI = 0.075 – 0.085; phage susceptible: 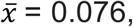 95%CI = 0.072 – 0.081; Figure 5). Phage resistant isolates show similar growth dynamics to susceptible isolates, except they maintain higher densities during the death phase at day 7.

**Figure 4.**
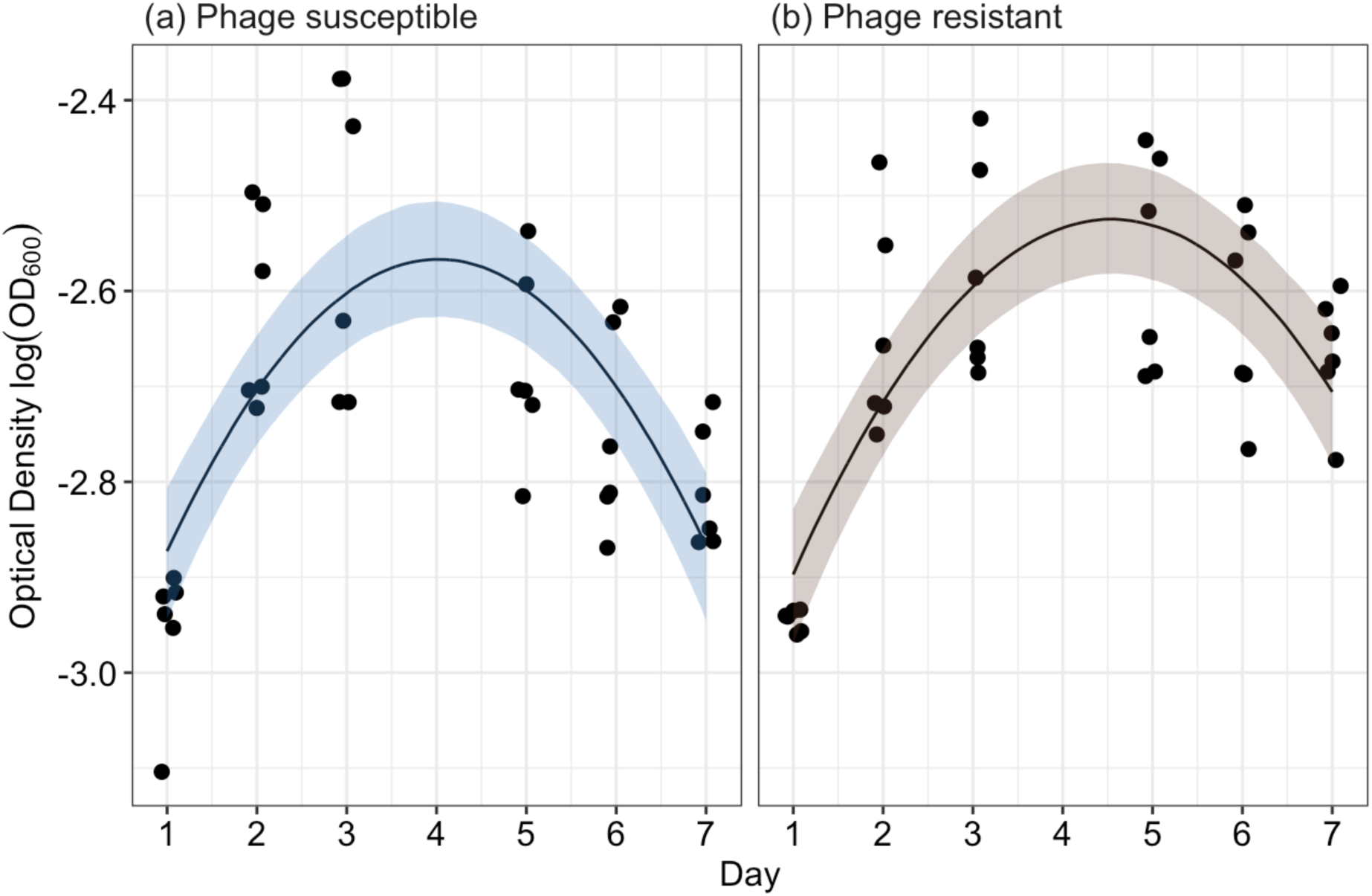
The seven-day growth curves of (a) phage susceptible and (b) phage resistant *Variovorax* populations that had evolved for seven weeks. Points represent individual replicate populations measured repeatedly through time. Solid lines represent the mean predictions, and shaded bands represent the 95% confidence intervals of the best fitted model.

**Figure 5.**
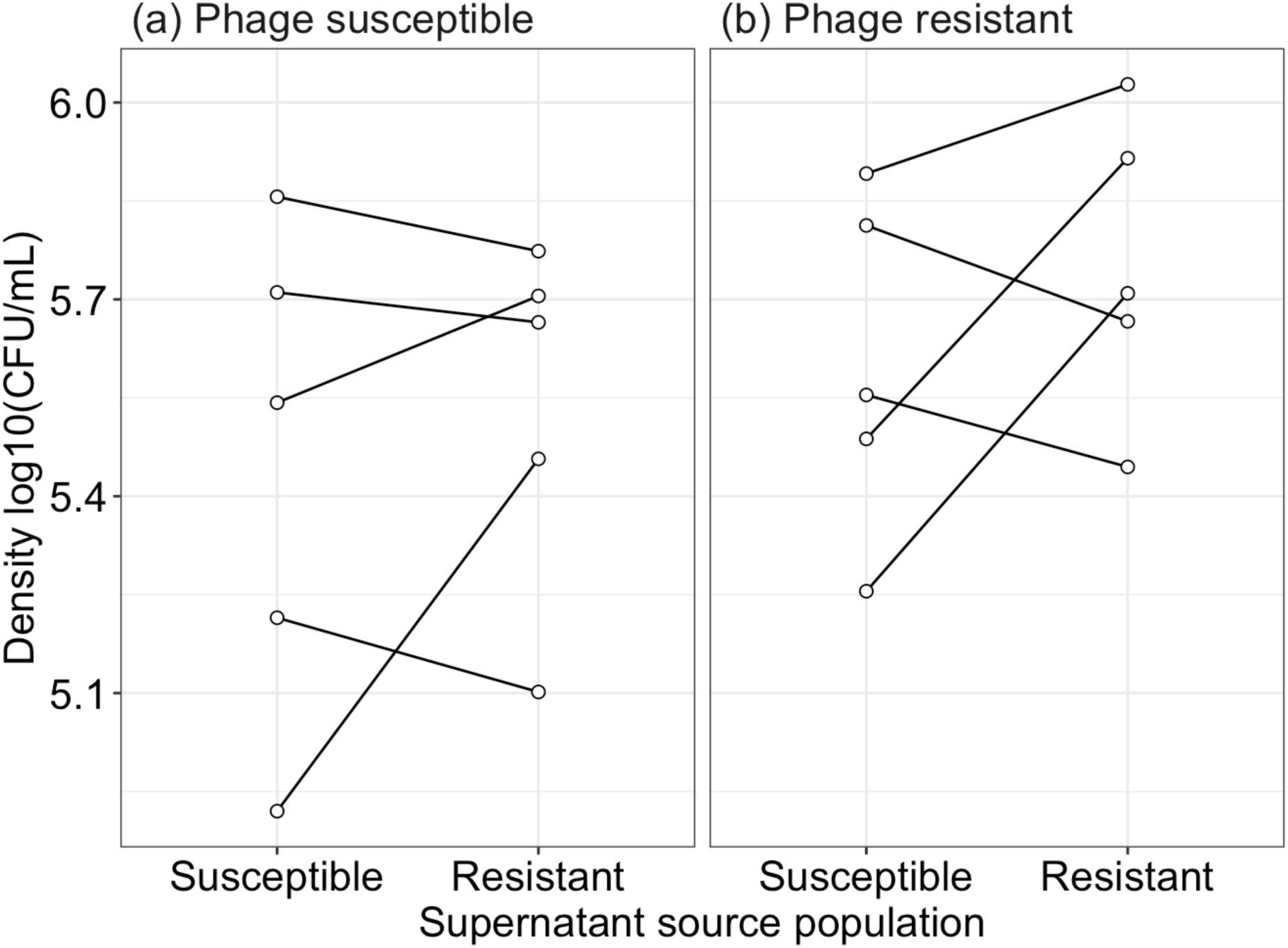
Density changes when populations which are (a) phage susceptible and (b) phage resistant, are grown in supernatant isolated from either population. E.g. phage susceptible growing in supernatant from phage susceptible populations (same evolutionary history) or phage susceptible growing in supernatant from phage resistant populations (different evolutionary history). Points represent densities of individual replicates while lines indicate the change when that same population is grown in a different supernatant.

### The supernatant of phage resistant populations is not less toxic

If bacterial populations can maintain a higher density during the stationary/death phase, this suggests that bacteria are potentially producing fewer toxic metabolites and/or are less susceptible to them. We investigated these possibilities by growing week seven populations in spent supernatants from either their own treatment or the opposite treatment. However, population density was non-significantly different between populations irrespective of prior phage exposure (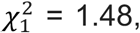 p = 0.224) or supernatant type (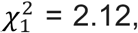 p = 0.145; interaction between effects: 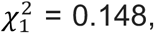 p = 0.7004; Figure 5).

### Indirect benefit of phage resistance impacts community structure

Considering that phage resistance increased *Variovorax* densities, we considered whether this had implications for the structure of a simple microbial community. We had previously cultured *Variovorax* with two other species, *Ochrobactrum* and *Pseudomonas*, from which it derives a fitness benefit to the other species’ detriment (Castledine *et al*. 2024b). As such, we predicted that phage resistance increasing *Variovorax*’s density may have negative implications for the other two species. To this end, we re-assembled communities with different *Variovorax* isolates with ancestral *Ochrobactrum* and *Pseudomonas*. Consistent with this prediction, the relative proportion of *Pseudomonas* in the community was significantly lower with phage resistant *Variovorax* (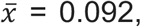 95%CI = 0.08 – 0.11) compared to ancestral (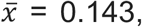 95%CI = 0.128 – 0.158; Tukey HSD comparing *Pseudomonas* proportion with ancestral vs resistant *Variovorax*: estimate = 0.495, z-ratio = 5.02, p < 0.001) or phage susceptible *Variovorax* (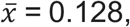 95%CI = 0.114 – 0.143; ANOVA comparing models with and without interaction between *Variovorax* phage resistance and species identity: 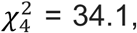 p < 0.001; Tukey HSD comparing *Pseudomonas* proportion with resistant vs susceptible *Variovorax*: estimate = -0.367, z-ratio = -3.64, p < 0.001; Figure 6). *Pseudomonas*’s relative proportion was non-significantly different when cultured with ancestral or phage susceptible *Variovorax* (Tukey HSD: estimate = 0.129, z-ratio = 1.403, p = 0.339). Neither *Ochrobactrum*’s or *Variovorax’*s relative proportion was significantly affected by phage resistance in *Variovorax* (Tukey HSDs > 0.05, Table S1; Figure 6).

**Figure 6.**
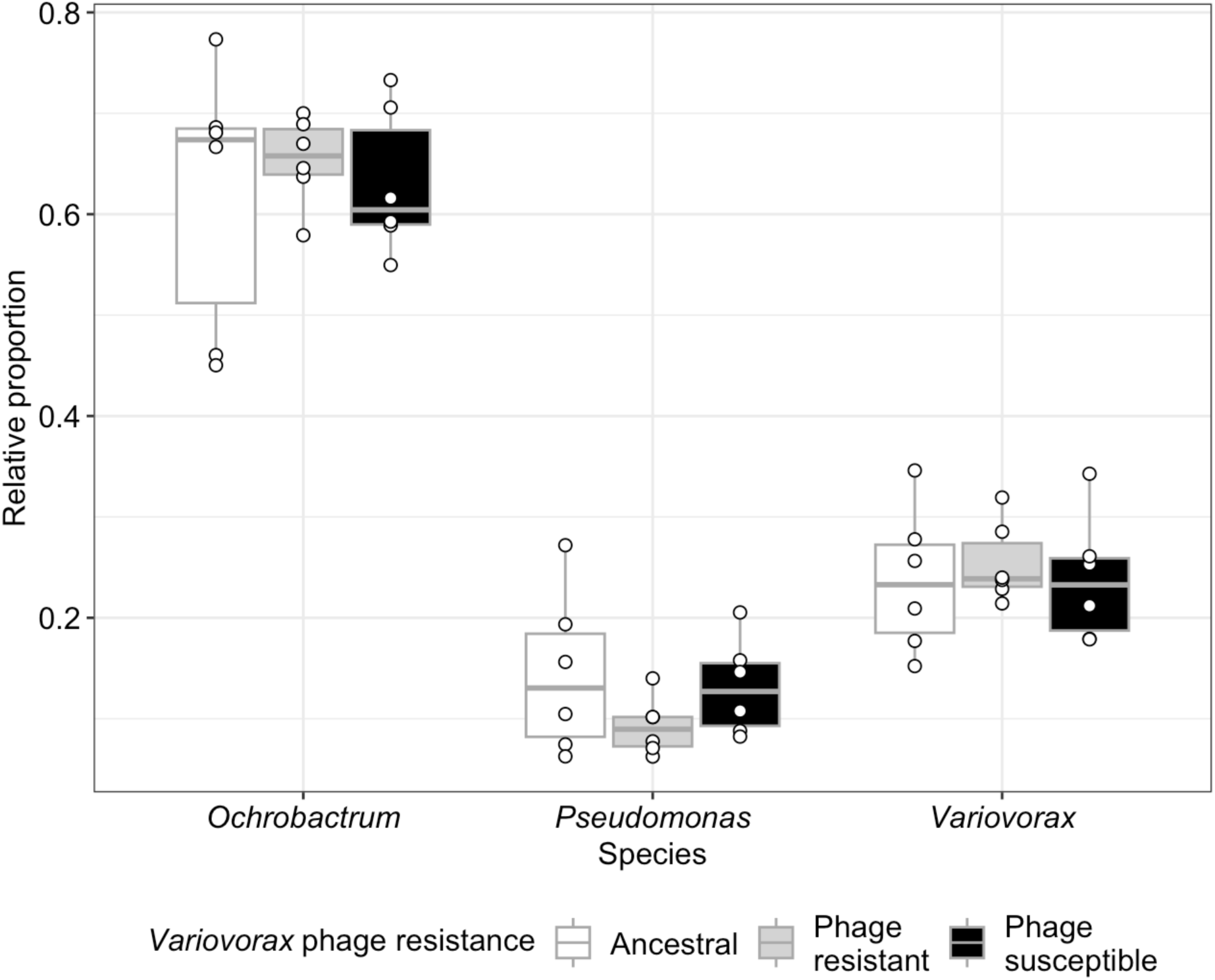
The relative proportion of each species in replicate communities when cultured with ancestral, phage resistant or phage susceptible *Variovorax* isolates. Points represent individual treatment replicates. Tops and bottoms of the bars represent the 75th and 25th percentiles of the data, white lines indicate the medians, and whiskers extend from their respective hinge to the smallest or largest value no further than 1.5× the interquartile range.

## Discussion

Bacteriophages have been previously shown to positively influence bacteria densities, but only when propagated in monoculture and the mechanism was not determined (Gómez and Buckling 2011). Here, we showed that evolved resistance to phage changed the growth curves of *Variovorax* populations resulting in higher densities during the death phase. High levels of resistance, and the associated positive effect on density, was observed for several weeks past phage extinction within monocultures (no other bacterial species present) and in a community context. That this effect was consistent in both the presence and absence of other species is particularly interesting as the growth of this species is enhanced by the presence of the other two species (Castledine, Padfield and Buckling 2020). As such, irrespective of the metabolic state the bacteria are in (feeding on the media or on supernatant), the phage had a positive effect on *Variovorax* density. This effect had consequences for community structure with phage resistance in *Variovorax* suppressing *Pseudomonas* in communities. These results demonstrate that phage resistance can give some species a relative advantage against others in community contexts, contrary to the prevailing wisdom of phage community ecology (Winter *et al*. 2010).

Phage resistance did not have a significant effect on exponential growth rate but did affect the death phase, with phage resistant populations maintaining a higher density. Although we did not find evidence that this was linked to a change in metabolites produced after one week, it may reflect a broader ability of the bacteria being able to cope in stressful conditions. *Variovorax* is a metabolically plastic species and is capable of being lithoautotrophic or chemoorganotrophic, including the metabolism of toxic or complex compounds (Satola, Wübbeler and Steinbüchel 2013). A lack of convergent evolution meant we could not link genotype to phenotype in this study. Clones had few genetic changes, but fitness benefits from phage-exposure were not linked to a single phage resistance mutation, although 2/3 clones had a missense mutation in the same protein. If this mutation and protein was the main mechanistic way phage-resistance increased fitness, we would expect it to also be present in the population sequencing, but no genetic changes in that locus were found. Instead, population sequencing revealed multiple mutations emerging and several reached high frequencies (>50%) in 5/6 phage treatment populations, although with a lack of convergent evolution. All high frequency genetic variants were in hypothetical proteins, making mechanistic inferences impossible within this project. Interestingly, only two no-phage culture had a unique mutation that reached high frequency (>50%) compared to 5/6 phage cultures, suggesting phage presence increased the likelihood of variants reaching a high frequency. Different mutations can confer the same phenotypic outcome, including phage resistance (Wright *et al*. 2018, 2019; Debray, De Luna and Koskella 2022), which is likely to be the case in this study.

While resistance was clearly beneficial in the presence and absence of phage, reaching and being maintained at very high frequencies, we did not find any phage resistance emerge in phage-free controls across eight weeks. This was presumably because there was no selective benefit of phage resistant mutants competing against phage sensitive clones in the absence of phage, and hence resistant mutations would not have increased in frequency. Nevertheless, we may expect this resistance-linked growth phenotype to be favoured in highly structured metapopulations, where limited dispersal can result in different patches seeded by single clones with the most productive patches (i.e. those seeded with resistant clones) producing more individuals to colonise new patches (Griffin, West and Buckling 2004). The resistance mutations may also allow populations to subsequently explore alternative evolutionary trajectories in the longer term (Salathé and Soyer 2008).

Importantly, phage resistance increasing *Variovorax* densities shifted community structure, resulting in lower proportions of *Pseudomonas*. *Variovorax* has been shown in previous work to benefit from the presence of *Pseudomonas* to *Pseudomonas*’ detriment (Castledine *et al*. 2024b). Increasing densities of phage resistant *Variovorax* therefore likely resulted in increased costs experienced by *Pseudomonas* in coculture. Phage lysis and phage resistance are typically assumed to be most costly to dominant community members (Winter *et al*. 2010). However, many examples exist of non-costly phage resistance and species dominating communities with high levels of phage resistance (Zhao *et al*. 2013; Castledine and Buckling 2024). Consequently, our results in-part support the king-of-the-mountain’ hypothesis in which dominant community members are more able to evolve resistance owing to greater population sizes (Zhao *et al*. 2013; Giovannoni 2017) – albeit in this case likely by mutation than horizontal-gene-transfer (starting population isogenic). Synonymously, the ‘king-of-the-mountain’ hypothesis also relies on a low trade-off in defence and competition (Thingstad *et al*. 2014), evident in *Variovorax*.

Few examples exist of phages improving bacteria densities (Gómez and Buckling 2011), suggesting this observation may not be ubiquitous. However, much research has heavily focussed on model strains of bacteria and phage which are unlikely to be generalisable to all bacteria and phages (Castledine and Buckling 2024; Castledine *et al*. 2024a). Furthermore, different mechanisms of phage resistance have the potential to improve bacteria densities in specific contexts. For example, phages can select for increased biofilm production (Scanlan and Buckling 2012; Castledine *et al*. 2022) and persister populations (Fernández-García *et al*. 2023) which can further confer resistance to stressors such as antibiotics and immune cells (Crabbé *et al*. 2019; Niu, Gu and Zhang 2024). If phages can improve the density of soil bacteria, such as by driving host diversification (Brockhurst, Rainey and Buckling 2004), this may have positive impacts on carbon cycling and crop yield (Dy, Rigano and Fineran 2018). *Variovorax* species, for instance, are symbionts to plants and are found in the rhizosphere where they can improve plant growth (Satola, Wübbeler and Steinbüchel 2013). Alternatively, if phages are used therapeutically (Oechslin 2018; Abedon 2019) then density enhancement of target bacterial pathogens is clearly detrimental. Work considering wider bacteria and phage pairs, beyond model systems, will allow greater generalisations to be made and understanding of how phages can affect their hosts in the short and long-term.

The influence of phages on bacterial populations are highly diverse, making predictions and generalisations challenging from model systems to phages in nature. Our work highlights the surprising outcomes that can arise from bacteria-phage interactions, and that the simple assumption that lytic phages negatively affect host densities may not always be true. Wider research analysing the effects of different phages on host bacteria, both during and after phage extinction would give important insight into how phage therapy may affect pathogens during infection and how phages operate in nature.

## Author contributions

MC and AB conceived and designed the study. Experiments conducted by MC. DNA extractions for population sequencing conducted by RL. Bioinformatics conducted by DP and MC. Data analysis conducted by MC. All authors contributed to the writing of the manuscript.

## Competing Interests

We have no competing interests

## Acknowledgements

We thank Stineke van Houte and Michael Brockhurst for helpful feedback on a manuscript draft. This work was funded by grant no. MR/N0137941/1 for the Great West 4 BIOMED Medical Research Council Doctoral Training Partnership, awarded to the Universities of Bath, Bristol, Cardiff, and Exeter from the Medical Research Council/UK Research and Innovation, awarded to MC. This work was supported by NERC awards NE/V012347/1 and NE/S000771/1 awarded to AB. DP is funded by a NERC independent research fellowship (NE/W008890/1).

## Data availability

All data and R code used in the analysis are available on GitHub (https://github.com/mcastledine96/Variovorax_phage_resistance_increases_density_ 2024). Assemblies and sequencing data are available from ENA (Project Accession PRJEB74093).

## Supplementary Information

## Supplementary results: pool sequencing

In total, we found 470 genetic variants, with 239 shared between resistant and susceptible populations. Phage resistant populations had 117 unique variants (within 60 genes) while phage susceptible populations had 114 unique variants (within 64 genes). Most variants were found at low frequency (96.8% of variants at <50% frequency) with resistant and susceptible populations having on average 147 variants. 5/6 phage resistant populations had at least one unique genetic variant at high frequency (>50%) compared to only 2/6 of the phage susceptible populations, suggesting specific mutations were selected by phage.

While we did not observe mutations in 01575 (mutation found in 2/3 resistant clones), mutations were found in 01584 (mutation found in 1/3 resistant clones) in 4/6 phage resistant populations (92.7%, 68.2%, 12.3%, 9.6%). Mutations selected to high frequency were also found in *prpR* (69.3% in 1 population; transcription regulator gene) and in five other hypothetical proteins or intergenic regions (Figure S1). The lack of convergence suggests phage resistance may be maintained by several different mechanisms that have the same phenotypic effects (resistance and higher population density). As a result, we did not find phage presence significantly differed in genetic composition (PERMANOVA: F_1,10_ = 0.889, p = 0.81, Figure S2) or distance from the ancestral genome (ANOVA: F_1,10_ = 0.547, p = 0.477).

**Figure S1.**
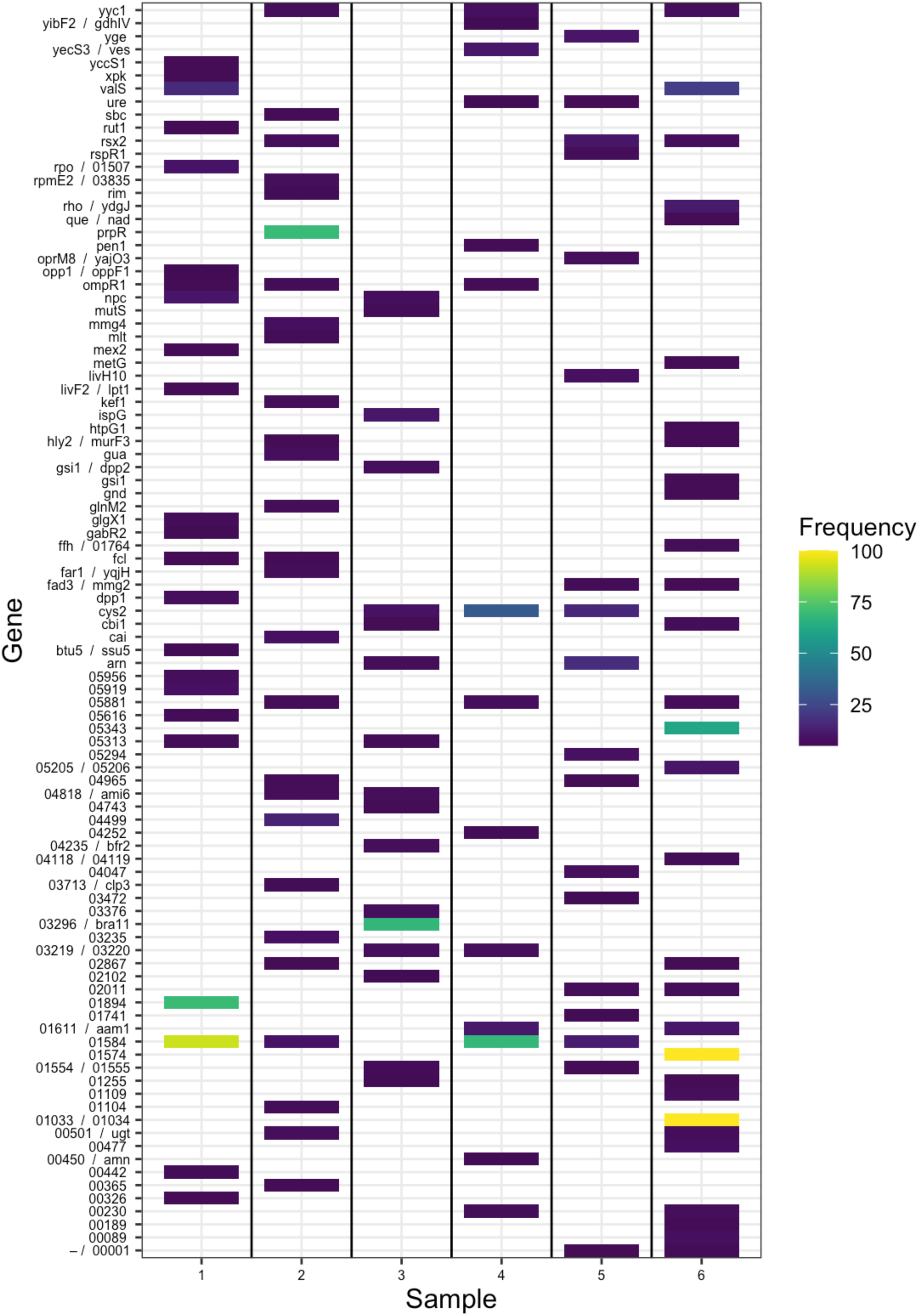
The frequency of genetic variants found in phage resistant but not phage susceptible populations. Where multiple variants are found in the same gene, the average has been presented. Each replicate population is identified on the x-axis. Genes of known function are named on the y-axis while hypothetical proteins are given a numerical denomination. Gene names separated by a forward slash indicate mutations in intergenic regions. Samples indicate each of the six replicate populations that evolved in the presence of phage.

**Figure S2.**
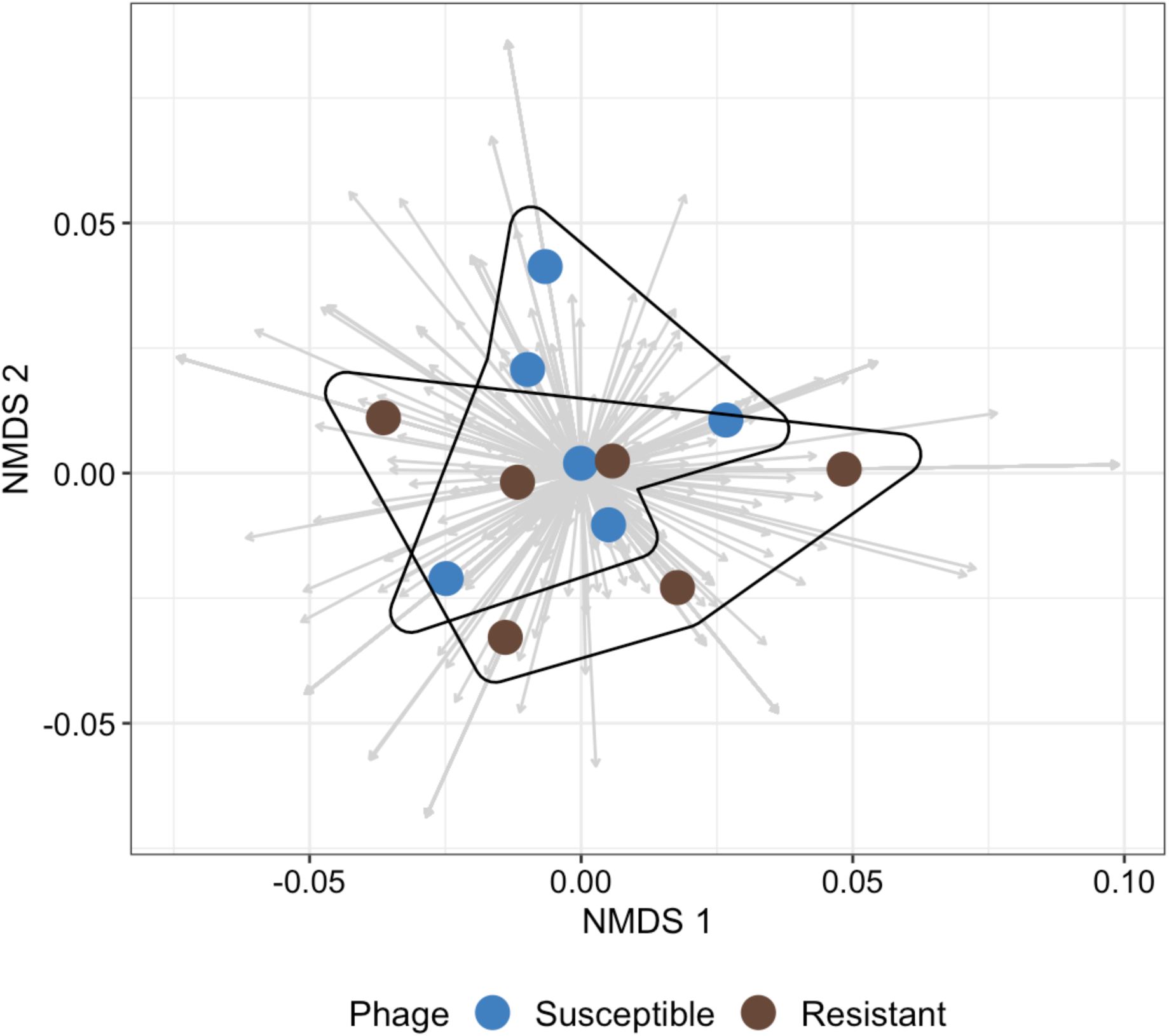
The divergence of each treatment group as indicated by Euclidean distances of each gene (arrows) from the ancestral genome. Points represent individual replicates sequenced from phage resistant or susceptible treatment groups.

## Supplementary Table

**Table S1.** Tukey HSD comparisons comparing the relative proportion of *Ochrobactrum* (O), *Pseudomonas* (P) and *Variovorax* (V) when the *Variovorax* strain is the ancestral type, phage resistant or susceptible. P-values are adjusted by the tukey method for comparing a family of three estimates.

| Species | Contrast | Estimate | z-ratio | p-value |
| --- | --- | --- | --- | --- |
| O | Ancestral - Resistant | -0.146 | -2.252 | 0.063 |
| O | Ancestral - Susceptible | -0.049 | -0.763 | 0.726 |
| O | Resistant - Susceptible | 0.096 | 1.483 | 0.299 |
| P | Ancestral - Resistant | 0.495 | 5.018 | <0.001 |
| P | Ancestral - Susceptible | 0.129 | 1.403 | 0.339 |
| P | Resistant - Susceptible | -0.367 | -3.637 | 0.001 |
| V | Ancestral - Resistant | -0.091 | -1.257 | 0.42 |
| V | Ancestral - Susceptible | -0.019 | -0.259 | 0.964 |
| V | Resistant - Susceptible | 0.072 | 0.995 | 0.58 |

## Notes

### Competing Interest Statement

The authors have declared no competing interest.

### Summary of Updates

Edits have been made to discussion of results following journal feedback.

